# NADPH oxidase promotes glioblastoma radiation resistance in a PTEN-dependent manner

**DOI:** 10.1101/2022.06.16.496502

**Authors:** Kirsten Ludwig, Janel E. Le Belle, Sree Deepthi Muthukrishnan, Jantzen Sperry, Michael Condro, Erina Vlashi, Frank Pajonk, Harley I. Kornblum

## Abstract

**Aims:** The goal of this study was to determine whether NADPH oxidase (NOX)-produced reactive oxygen species enhances brain tumor growth of glioblastoma (GBM) under hypoxic conditions and during radiation treatment.

**Results:** Exogenous ROS promoted brain tumor growth in gliomasphere cultures that expressed functional PTEN, but not in tumors that were PTEN deficient. Hypoxia induced the production of endogenous cytoplasmic ROS and tumor cell growth via activation of NOX. NOX activation resulted in oxidation of PTEN and downstream Akt activation. Radiation also promoted ROS production via NOX which, in turn, resulted in cellular protection that could be abrogated by knockdown of the key NOX component, p22. Knockdown of p22 also inhibited tumor growth and enhanced the efficacy of radiation in PTEN-expressing GBM cells.

**Innovation:** While other studies have implicated NOX function in GBM models, these studies demonstrate NOX activation and function under physiological hypoxia and following radiation in GBM, two conditions that are seen in patients. NOX plays an important role in a PTEN-expressing GBM model system, but not in PTEN-non-functional systems and provide a potential, patient-specific therapeutic opportunity.

**Conclusions:** This study provides a strong basis for pursuing NOX inhibition in PTEN-expressing GBM cells as a possible adjunct to radiation therapy.

## INTRODUCTION

Glioblastoma (GBM) is a devastating disease, with a median survival of only 12-15 months. While the current standard of care of maximal surgical resection followed by radiation and chemotherapy with temozolomide, prolongs life, tumors virtually always recur (Vilar et al., 2022). The mechanisms of this recurrence are multifaceted and may include cellular plasticity that is induced by chemo and radiotherapy, as well as stem-like cells within tumors that are inherently resistant to these treatments, the so-called glioma stem cell (GSC) (Ludwig and Kornblum, 2017).

Although the cell of origin of GSC is unclear and potentially varied, these cells have some of the properties of neural stem cells (NSC), including their ability to self-renew and grow as gliomaspheres in a relatively simple, defined medium under the mitogenic control of added epidermal growth factor (EGF) and basic fibroblast growth factor (FGF) (Hemmati et al., 2003). Prior work in NSC biology has proven a role for the activation of the PI3K/Akt pathway in normal NSC proliferation, including self-renewal, and maintenance of neurogenesis (Groszer et al., 2006, 2001). One means by which exogenous factors can influence this pathway is through the activation of the NADPH oxidase (NOX) complex. NOX is found in abundance in many cell types, and its activation in neutrophils results in killing of the engulfed bacteria (Vermot et al., 2021). In neural stem cells, as well as a variety of cancers, NOX is activated by growth factors and other means, to oxidatively inactivate the PTEN protein, resulting in enhanced Akt and mTOR activity (Le Belle et al., 2014, 2011).

The GBM microenvironment is primed to activate NOX, given the common occurrence of hypoxia and the presence of several growth factors and cytokines that are often associated with GBM (Rodriguez et al., 2022). Furthermore, radiation itself has been demonstrated to activate NOX (Mortezaee et al., 2019). Therefore, we hypothesized that NOX activation could play an important role in GBM growth and radiation resistance. Indeed, a prior report using model systems has provided some evidence for the former (Cao et al., 2022; Su et al., 2021). Here, we examined the role of NOX activation in radiated and non-radiated GBM cells and found that both hypoxia and radiation induced NOX activity to inactivate PTEN, enhance Akt phosphorylation and increase cell numbers (Graphical Abstract; Fig. 1). Reducing NOX activity resulted in slower tumor growth *in vivo* and enhanced radiosensitivity in PTEN expressing tumors. These findings suggest that inhibition of NOX activation is a potential therapeutic target in PTEN-functional GBM.

**Figure 1:**
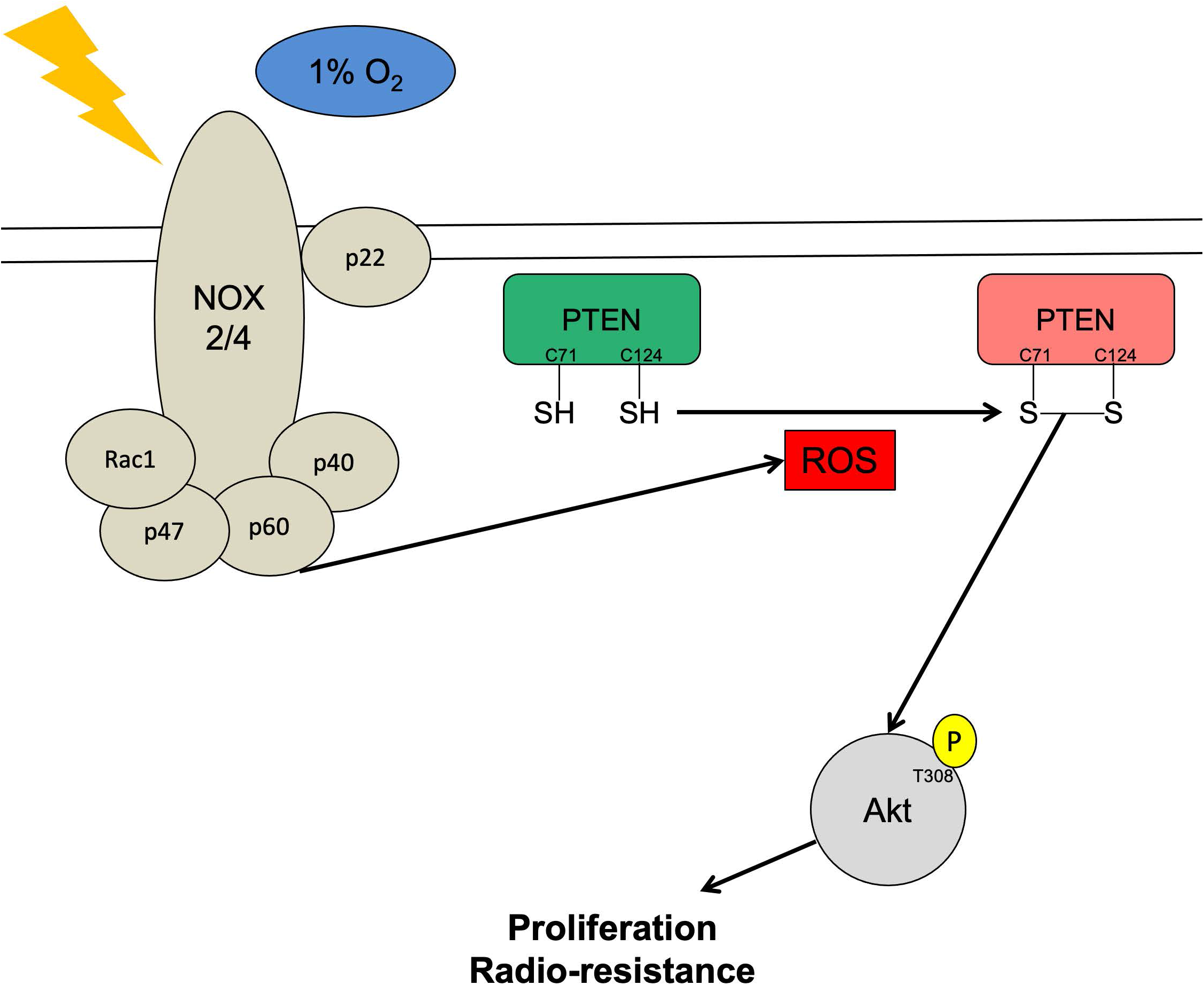
NOX-generated ROS promotes PTEN oxidation and inactivation to promote the Akt survival pathway following exposure to radiation or oxidation. A) Graphical representation of radiation or hypoxia activating NOX which raises ROS levels in the cytoplasm. Increased ROS levels oxidize and inactivate PTEN, which increases Akt signaling and promotes proliferation and resistance to radiation.

## RESULTS

Several studies, including those from our own lab, have shown that modestly elevated ROS levels are able to promote proliferation in cancer and some non-transformed cells, including neural progenitors (Le Belle et al., 2014, 2011). In primary GBM lines, we found that cells with higher endogenous ROS levels were significantly correlated with a faster proliferation rate (Fig. 2A), suggesting that ROS may promote proliferation in patient derived GBM lines. Data from our lab has previously shown that ROS-induced proliferation in neural stem cells occurs in a PI3K/Akt dependent manner (Le Belle et al., 2011). Similarly, in GBM, we found that treatment of cells that expressed functional PTEN protein with hydrogen peroxide (H_2_O_2_) (concentrations were and time points were chosen following a time course and dose dependent response [data not shown]) resulted in increased cell number (Fig. 2B), while those without functional PTEN protein did not respond to H_2_O_2_ treatment, even though treatment resulted in elevated ROS levels in PTEN-functional and non-functional cells (Fig. 2C). Conversely, a reduction in ROS levels, through treatment with the antioxidant N-acetylcysteine (NAC), resulted in decreased cell number in PTEN functional cells with no change in PTEN non-functional cells. Because PTEN function may not be the only difference between these lines, we directly examined the role of PTEN in this process by lentiviral knockdown of PTEN. This knockdown prevented ROS-induced increase in cell number in all 3 PTEN functional lines used (Fig. 2D). Efficacy was observed with the two different PTEN shRNAs that exhibited knockdown of the PTEN protein, while the two constructs exhibiting no significant knockdown produced no functional effect (Supp. Fig. 1A-C). Similarly, a decrease in ROS levels following NAC treatment resulted in decreased cell number in control treated cells, but not in the PTEN KD cells. No difference in ROS levels were found between the shCON and shPTEN groups (Fig. 2E) suggesting the changes are directly due to PTEN. Taken together, these data suggest that ROS increases cell number in patient derived GBM cells in a PTEN-dependent manner.

**Figure 2:**
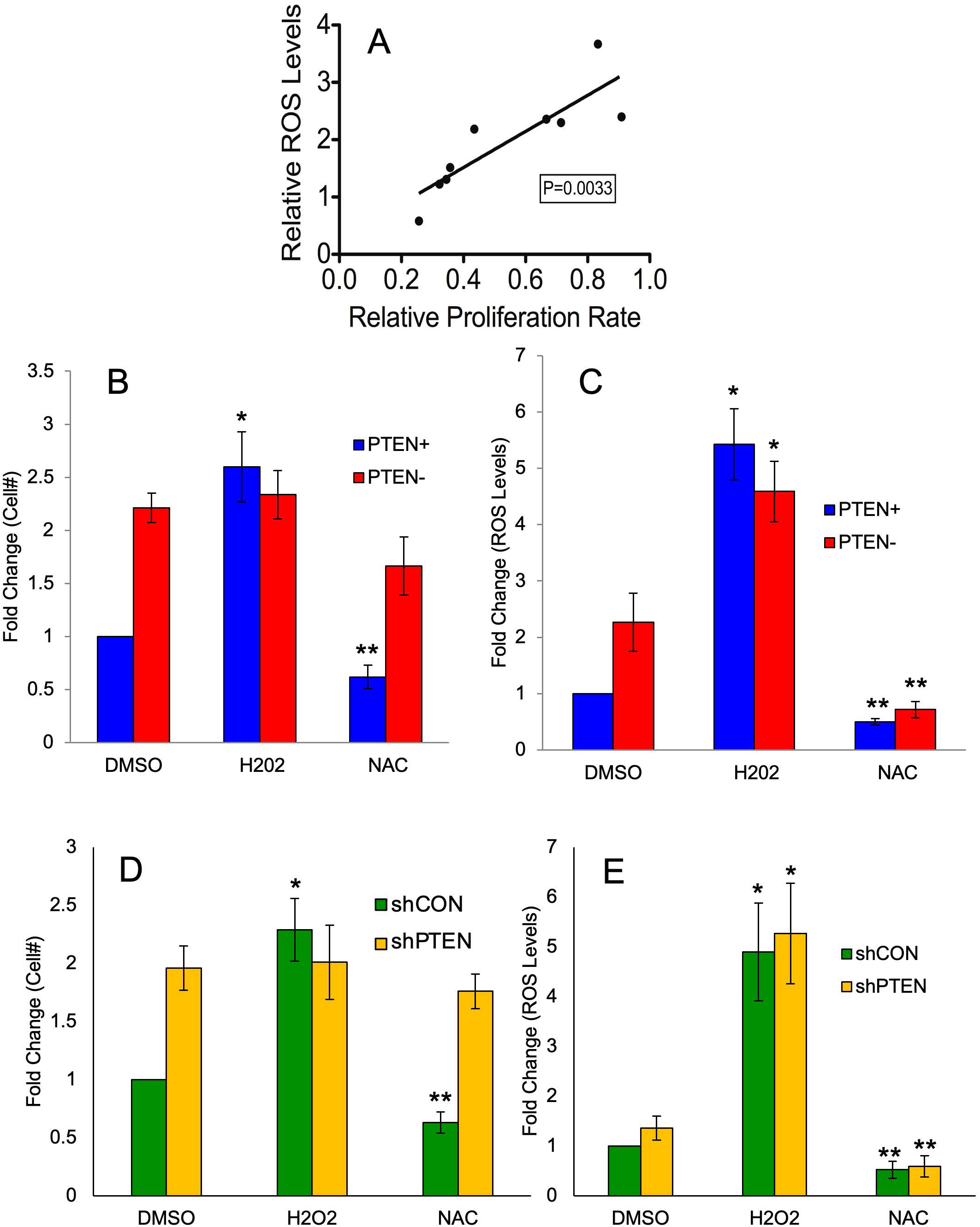
ROS induced proliferation requires PTEN. A) Correlation of endogenous ROS levels with relative proliferation rate in HK157, HK217, HK229, HK296, HK350, HK351, HK374, HK382, and HK 393 cells. B) Changes in cell number were determined with a CCK-8 assay following treatment with 10uM H_2_O_2_ or 1mM N-acetylcysteine (NAC) for 5 days in three PTEN functional lines (HK157, HK339, and HK374) and three PTEN non-functional lines (HK217, HK229, and HK296). Average values of all three lines were used to calculate data. n>3, *=*p*-value<0.01 as compared to DMSO, **=*p*-value<0.05 as compared to DMSO. C) Changes in ROS levels were determined with CellROX Green following treatment with 10uM H_2_O_2_ or 1mM N-acetylcysteine (NAC) for 5 days in three PTEN functional lines (HK157, HK339, and HK374) and three PTEN non-functional lines (HK217, HK229, and HK296). Average values of all three lines were used to calculate data. n>3, *=*p*-value<0.01 as compared to DMSO, **=*p*-value<0.05 as compared to DMSO. D) Changes in cell number were determined with a CCK-8 assay following treatment with 10uM H_2_O_2_ or 1mM N-acetylcysteine (NAC) for 5 days in three PTEN functional lines (HK157, HK339, and HK374) with either shCON or shPTEN. n>3, *=*p*-value<0.01 as compared to DMSO, **=*p*-value<0.05 as compared to DMSO. E) Changes in ROS levels were determined with CellROX Green following treatment with 10uM H_2_O_2_ or 1mM N-acetylcysteine (NAC) for 5 days in three PTEN functional lines (HK157, HK339, and HK374) with either shCON or shPTEN. Average values of all three lines were used to calculate data. n>3, *=*p*-value<0.01 as compared to DMSO, **=*p*-value<0.05 as compared to DMSO.

Increased ROS levels, like addition of H_2_O_2_, have been shown to oxidize PTEN resulting in its inactivation due to the creation of covalent disulfide bonds between cystine residues (Zhang et al., 2020). However, this has rarely been documented in cells under biologically relevant conditions (like low oxygen). Cancer cells are often found in low oxygen environments, which has been shown to paradoxically increase cytoplasmic ROS levels (Rodriguez et al., 2022). GBM cells grown under low oxygen conditions (1% O_2_) resulted in PTEN oxidation and increased Akt phosphorylation (Fig. 3A) suggesting that PTEN was inactivated under low oxygen conditions. PTEN oxidation under low oxygen was also associated with increased cell number (Fig. 3B) and increased ROS (Fig. 3C) (Cells were acclimated to low oxygen conditions for two weeks prior to experiment). Although PTEN knockdown cells also demonstrated increased ROS levels under hypoxia, this did not result in enhanced cell numbers, suggesting that PTEN inactivation via oxidation may be required for this effect in GBM cells under hypoxic conditions.

**Figure 3:**
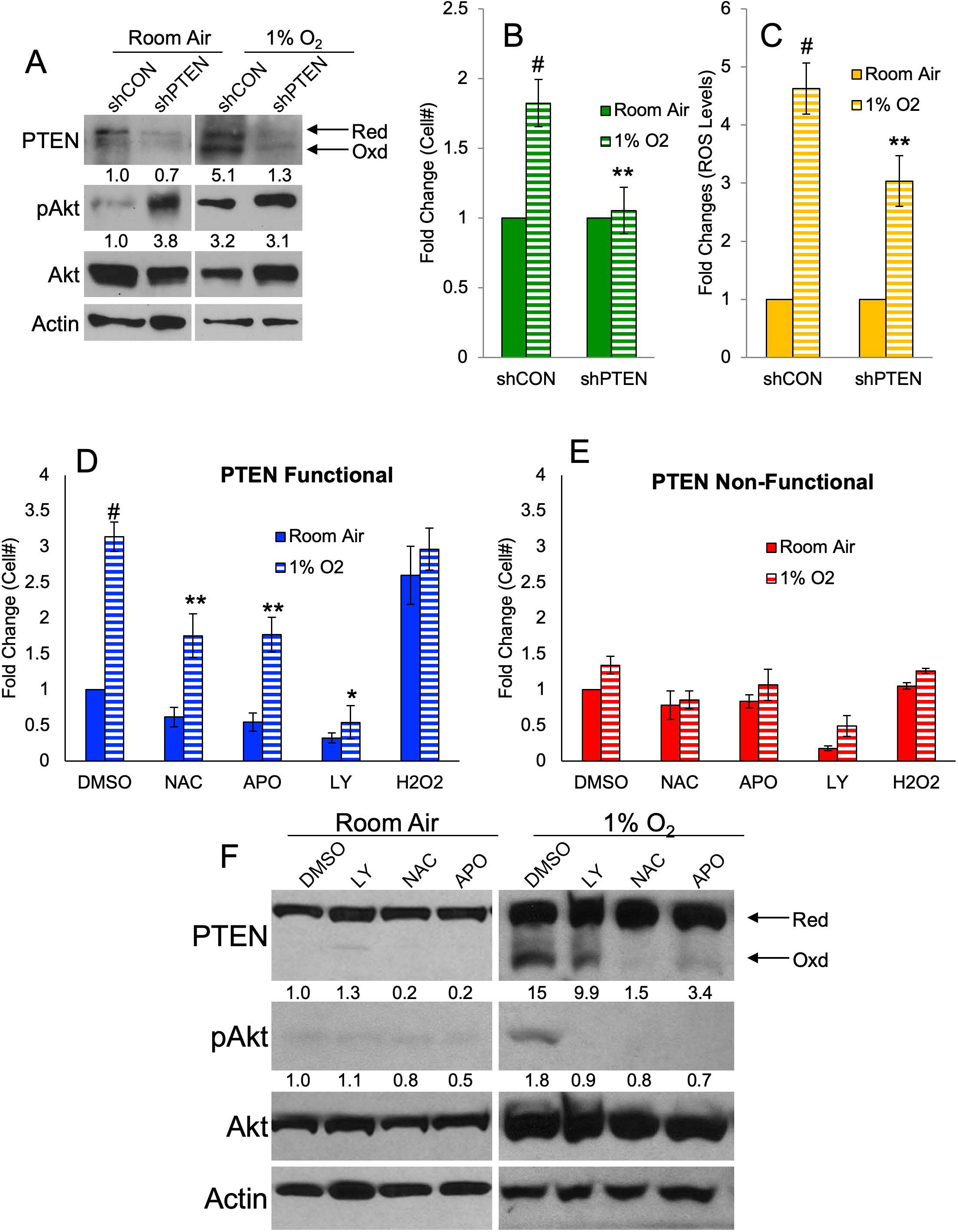
Hypoxia promotes PTEN oxidation and ROS-induced proliferation. A) Western blot of shCON and shPTEN cells (HK157) grown under room air or 1% O_2_ for 5 days indicating PTEN oxidation and total and phospho-Akt with actin used as a loading control. PTEN quantitation is of oxidized PTEN. Image is representative n>3 B) Changes in cell number were determined with a CCK-8 assay in shCON or shPTEN cells (HK157) grown under room air or 1% O_2_ for 5 days. n>3, #=*p*-value<0.01 as compared to room air, *=*p*-value<0.01 as compared to DMSO, **=*p*-value<0.05 as compared to shCON. C) Changes in ROS levels were determined with CellROX Green in shCON or shPTEN cells (HK157) grown under room air or 1% O_2_ for 5 days. n>3, #=*p*-value<0.01 as compared to room air, *=*p*-value<0.01 as compared to DMSO, **=*p*-value<0.05 as compared to shCON. D) Changes in cell number were determined with a CCK-8 assay in a PTEN functional line (HK157) grown under room air or 1% O_2_ for 5 days and treated with NAC (1mM), APO (100uM), LY (20uM), or H_2_O2 (10uM). n>3, #=*p*-value<0.01 as compared to room air, **=*p*-value<0.05 as compared to DMSO, 1% 02, *=*p*-value<0.01 as compared to DMSO, 1% O_2_ E) Changes in cell number were determined with a CCK-8 assay in a PTEN non-functional line (HK217) grown under room air or 1% O_2_ for 5 days and treated with NAC (1mM), APO (100uM), LY (20uM), or H_2_O2 (10uM). n>3 F) Western blot of a PTEN functional line (HK157) grown under room air or 1% O_2_ for 5 days and treated with NAC (1mM), APO (100uM), or LY (20uM). PTEN oxidation and total and phospho-Akt with actin used as a loading control. PTEN quantitation is of oxidized PTEN. Image is representative n>3

To better understand this pathway, a PTEN functional line and a PTEN non-functional line were grown under hypoxic conditions and cell number (Fig. 3D, E) and ROS levels (Supp. Fig. 2A-B) were assessed. As expected, the PTEN functional line exhibited increased cell numbers when grown in hypoxic conditions, while the PTEN non-functional line did not show changes in cell numbers. Treatment of PTEN functional cells in hypoxic conditions with either NAC or the NADPH oxidase (NOX) inhibitor apocynin (APO) resulted in reduced cell numbers, suggesting that quenching of the ROS species was able to decrease the response and that the enhanced elevation of ROS could be partially due to NOX activation. Treatment of cells grown in room air with H_2_O_2_, resulted in elevated cell numbers. However, H_2_O_2_ treatment did not increase cell number when grown under hypoxic conditions, suggesting that hypoxia results in a level of ROS that maximally stimulates the cells. Finally, treatment with the PI3K inhibitor, LY294002, resulted in significant decrease in cell number independent of the hypoxic environment. The PTEN non-functional line, however, did not display significant changes in cell numbers when ROS levels were altered, but did show a decrease in cell number when the PI3K pathway was inhibited.

When PTEN functional cells were grown in low oxygen, this resulted in PTEN oxidation and an increase in Akt phosphorylation which was lost following the addition of NAC or APO (Fig. 3F). Importantly, LY treatment of hypoxic cells decreased Akt phosphorylation, but did not prevent PTEN oxidation. Treatment also decreased cell numbers but had no effect of ROS levels. This suggests that the ROS-induced proliferative response is dependent upon the oxidation and inactivation of PTEN resulting in an increase in the Akt pathway activation. Altogether, our data suggest that elevated ROS levels, whether via exogenous administration of H_2_O_2_ or induction of ROS by growth under hypoxic conditions, oxidatively inactivates PTEN, resulting in enhanced Akt activity, and the induction of GBM cell proliferation and/or survival. Apocynin is a purported inhibitor of NADPH oxidase (NOX), which is a major contributor to intracellular ROS production (Mortezaee et al., 2019), and so we hypothesized that the response involved the NOX complex.

NOX is a complex of proteins with several different isotypes to each component. NOX2 and p22 are localized to the cellular membrane. Upon activation of Rac1 the cytosolic components of NOX (p40, p47 and p60) translocate to the membrane bound complexes. This binding results in activation of NOX and generation of ROS. There are 5 different isotypes of NOX, as well as two isotypes of DUOX, all of which produce super oxide, which is quickly converted to hydrogen peroxide (Mortezaee et al., 2019). To determine if our cells express any of the components of this complex, qRT-PCR was performed on PTEN functional and non-functional lines (Fig. 4A). Components of the NOX complex were expressed regardless of the PTEN status, with NOX4 having the highest expression in both lines, which is consistent with sequencing data showing NOX2 and NOX4 with the highest expression in brain tissue (McKetney et al., 2019). NOX4 is unique among the family of proteins, as it does not require the cytosolic components to become activated, only membrane bound p22 (Mortezaee et al., 2019). GlioVis data (Bowman et al., 2017) shows that CYBA (NOX2), CYBB (p22) and NOX4 are all upregulated in glioblastoma as compared to non-tumor tissue (Fig. 4B). To directly address the role of NOX in this pathway, we targeted p22 with shRNA (Supp. Fig. 3A, B), as both NOX2 and NOX4 are dependent upon this protein to become activated. Knockdown of p22 in a PTEN functional line resulted in decreased cell number (Fig. 4C) and ROS levels (Fig. 4D) under room air conditions, as well as hypoxic conditions. Similar effects were observed with the three different shp22 constructs that resulted in 60% or more knockdown, while treatment with ineffective constructs had no biological effects (Supp. Fig 3A, C, D). It is important to note that APO and p22 knockdown were not able to completely protect cells from ROS production, as hypoxia most likely produced ROS through alternative mechanisms (Bekhet and Eid, 2021). Our data suggests that NOX is a major contributor to ROS formation, and likely to the ROS-indued proliferation, however it is not the sole source of ROS. Additionally, p22 knockdown had a greater effect than the NOX inhibitor, apocynin, most likely due to apocynin being quenched and used up by the cell, while p22 knockdown is constitutive. In addition, knockdown of p22 resulted in a decrease in PTEN oxidation and Akt phosphorylation (Fig. 4E) again implicating the role of the PI3K/AKT pathway in this process. Together, our data indicate that endogenous NOX induces ROS which then inactivate PTEN, resulting in elevated cell numbers under room air and hypoxic conditions, with the latter producing a greater activation of NOX.

**Figure 4:**
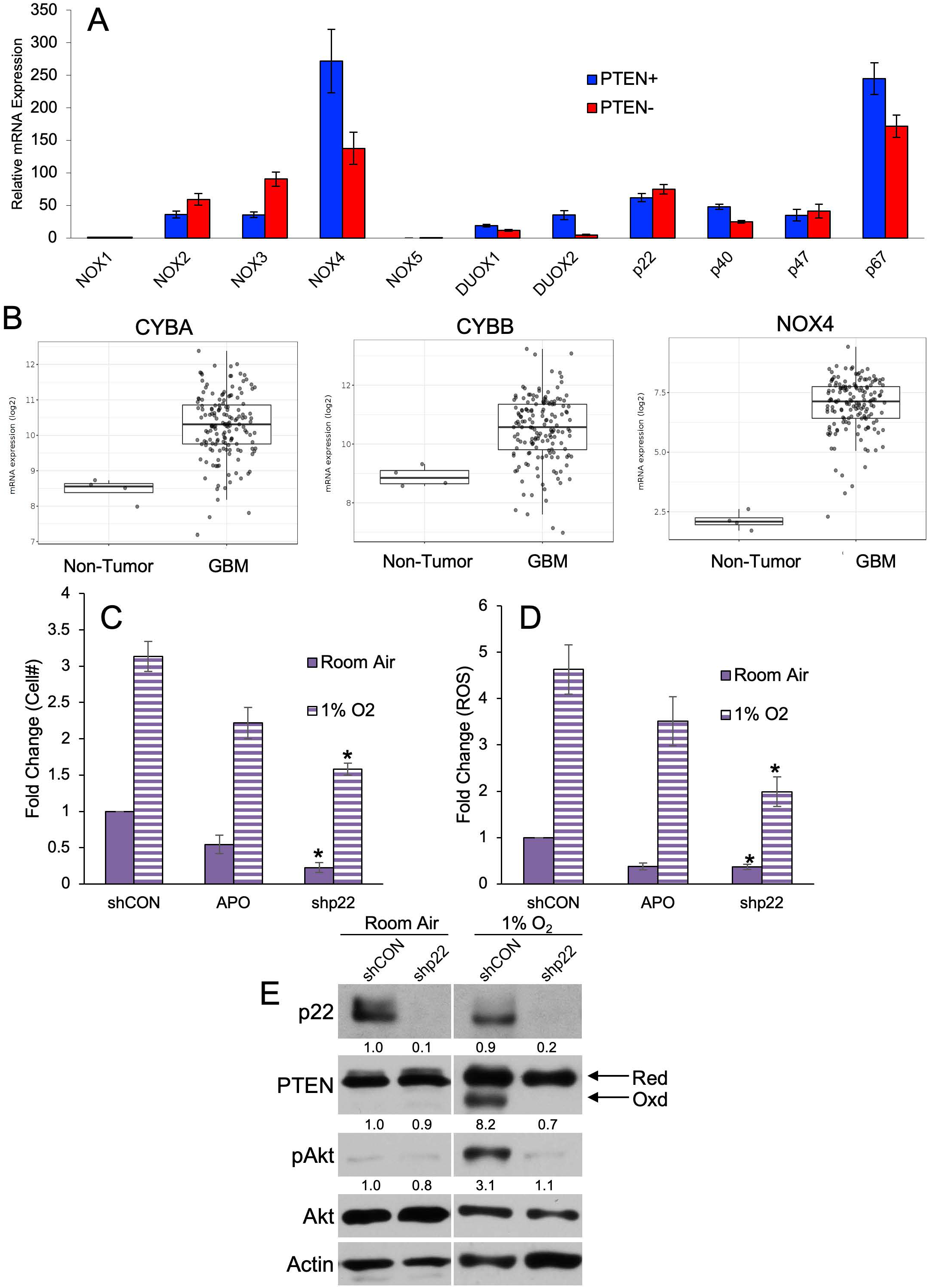
Hypoxia induces ROS and proliferation via NOX. A) mRNA levels of NOX components were assessed by q-RT-CPR in a PTEN functional (HK157) and non-functional (HK217) line under normal growth conditions. n=3 B) mRNA expression of NOX2 (CYBB), NOX4 and p22phox (CYBA) in tumor and GBM samples collected by TCGA. Data generated using GlioVis. C) Changes in cell number were determined with a CCK-8 assay in shCON or shp22 cells (HK157) grown under room air or 1% O_2_ or treated with APO (100uM) for 5 days. n>3, *=*p*-value<0.01 as compared to shCON D) Changes in ROS levels were determined with CellROX Green in shCON or shPTEN cells (HK157) grown under room air or 1% O_2_ or treated with APO (100uM) for 5 days. n>3, *=*p*-value<0.01 as compared to shCON E) Western blot of a PTEN functional line (HK157) grown under room air or 1% O_2_ for 5 days following KD of p22phox. PTEN oxidation and total and phospho-Akt with actin used as a loading control. PTEN quantitation is of oxidized PTEN. Image is representative n>3

Radiation, a mainstay of glioma treatment, results in elevated intracellular ROS, ultimately resulting in cell death and senescence (Chen et al., 2021). However, a subpopulation of glioma cells are refractory to radiation treatment and, in fact, demonstrate enhanced stem cell-like characteristics following radiation. These cells are the likely seeds of recurrence following radiation (Ludwig and Kornblum, 2017). The NOX complex of proteins have been shown to play a role in radio-resistance in a variety of tumor cells (Mortezaee et al., 2019), however the mechanism of how NOX promotes resistance to radiation is poorly understood. We hypothesized that our proposed NOX-PTEN axis may play a role in radiation resistance. We found that a single dose of radiation significantly increased the mRNA expression of NOX4 (Fig. 5A). Irradiating gliomaspheres resulted in enhanced total cellular ROS production and mitochondrial ROS (Fig. 5B). Knockdown of p22 resulted in diminished total ROS, but not mitochondrial ROS, indicating that radiation induces ROS via different mechanisms in these two compartments, with NOX activation being responsible for induction in the cytosolic compartment, the cellular compartment that houses the PTEN-Akt pathway. To further explore the effects of radiation-induced NOX, we examined γH2AX phosphorylation, a marker of DNA damage following a single dose (10 Gy) of radiation. As expected, irradiating shCON cells resulted in a greater than 2-fold increase in the percent of cells showing γH2AX phosphorylation, while knockdown of p22 significantly enhanced the number of positively stained cell, resulting in an over 3-fold increase following radiation (Fig. 5C). To better understand the role of PTEN loss in the protection of NOX-generated ROS from radiation, we compared cell numbers (Fig. 5E) and ROS levels (Fig. 5F) in PTEN functional and PTEN non-functional lines with p22 knockdown following radiation. We found that in both cell lines ROS levels were decreased with p22 knockdown before and after radiation. However, only the PTEN functional line exhibited a decrease in cell number following radiation in combination with p22 knockdown.

**Figure 5:**
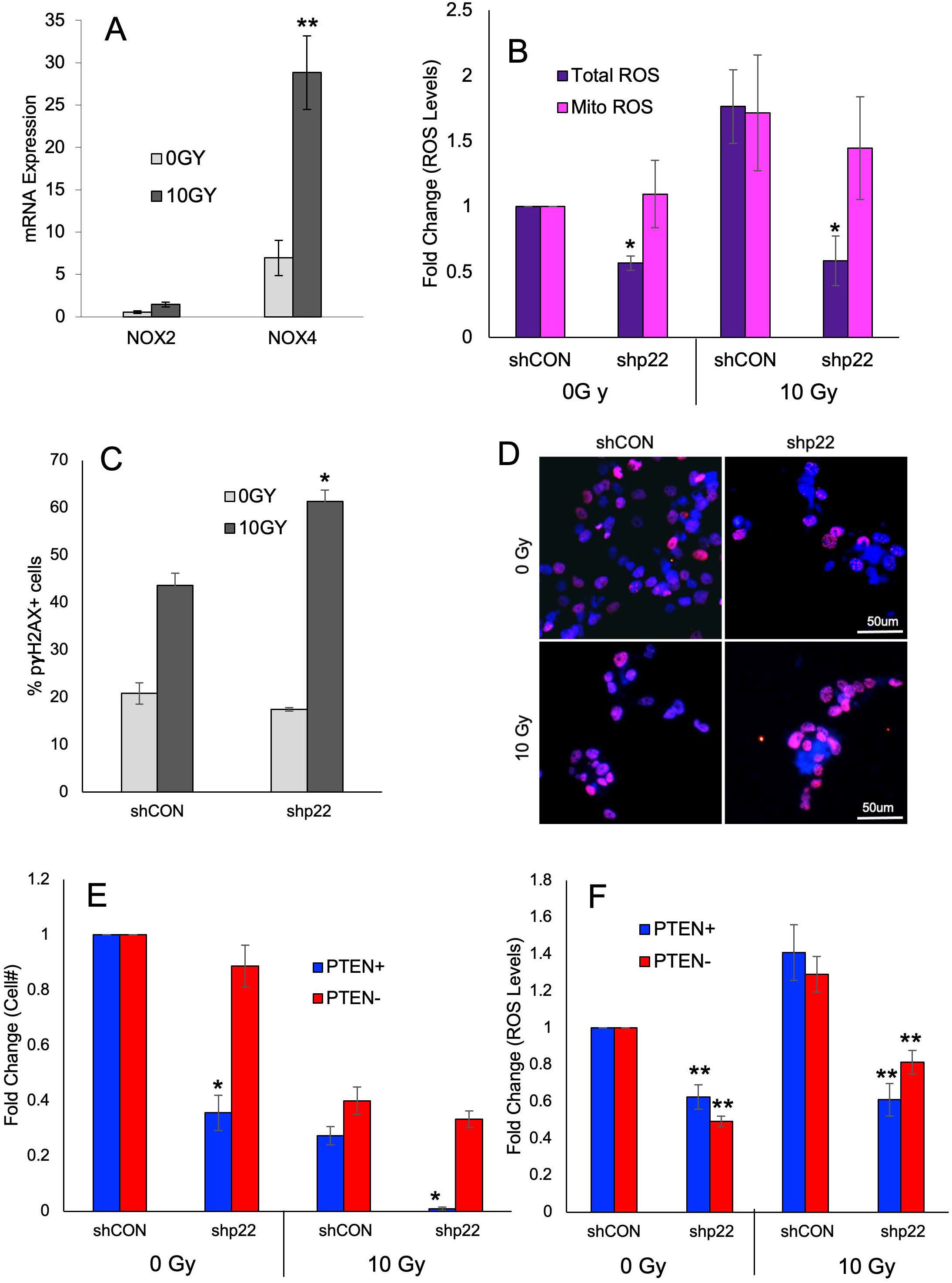
Radiation activates NOX to produce cytosolic ROS. A) mRNA levels of NOX2 and NOX4 were assessed by q-RT-CPR in a PTEN functional (HK157) 24 hours after exposure to 10 Gy. n>3, **=*p*-value<0.05 as compared to 0 Gy B) Changes in total ROS levels were determined with CellROX Green, and mitochondrial ROS with MitoSOX in shCON or shp22 cells (HK157) 30 min. after exposure to 0 Gy or 10 Gy. n>3, *=*p*-value<0.01 as compared to shCON C) pγH2AX levels were determined by immunocytochemistry in shCON or shp22 cells (HK408) 24 hours after radiation with 10 Gy. n=3, **=*p*-value<0.05 as compared to 0GY D) Representative images of shCON or shp22 cells (HK408) 24 hours after radiation with 10 Gy E) Changes in cell number were determined with a CCK-8 assay in shCON or shp22 cells (HK157 and HK217) 5 days after exposure to 0 Gy or 10 Gy. n>3, *=*p*-value<0.01 as compared to shCON F) Changes in ROS levels were determined with CellROX Green in shCON or shPTEN cells (HK157 and HK217) 5 days after exposure to 0 Gy or 10 Gy. n>3, **=*p*-value<0.05 as compared to shCON

To better define the role of PTEN in NOX-induced radiation resistance, we examined PTEN oxidation and Akt activation in PTEN functional and non-functional lines. In the PTEN functional line, PTEN was oxidized following a single dose of radiation which corresponded with an increase in Akt activation. This response was not seen in the PTEN non-functional line. However, knockdown of p22 rescued PTEN oxidation and prevented the increase in Akt activation (Fig. 6A). Our data suggest that NOX is activated following radiation to produce ROS which oxidizes and inactivates PTEN; this in turn increases Akt phosphorylation, which promotes proliferation and decreases the effect of radiation. However, PTEN non-functional cells already exhibit increased Akt activation, and hence do not display enhanced proliferation, suggesting that the presence of a functional PTEN is required for NOX inhibition to decrease radio-resistance. We found that knocking down PTEN in conjunction with p22 and radiation reversed the shp22 effect (Fig. 6B) even though ROS levels were decreased (Fig. 6C). Together, this supports our hypothesis that NOX is required for ROS generation following radiation, which oxidizes PTEN and increases the pro-proliferative signals promoting radiation resistance.

**Figure 6:**
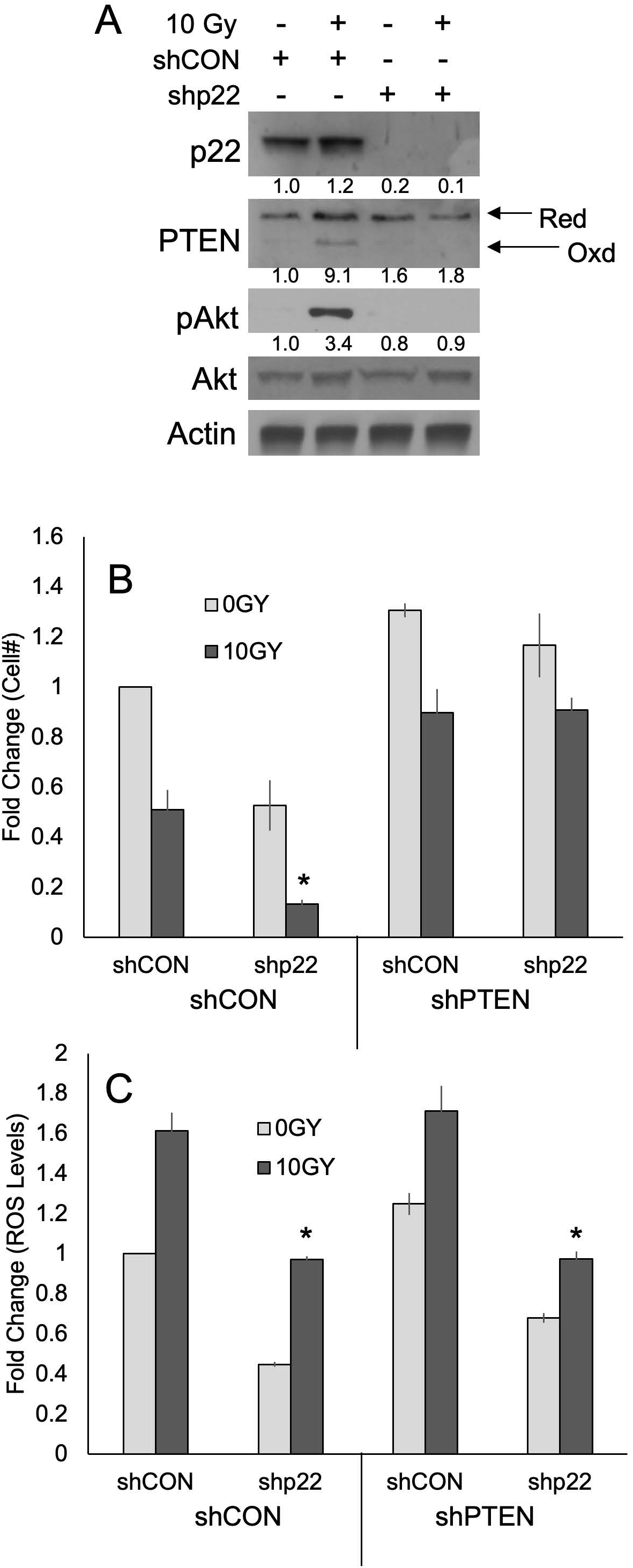
PTEN oxidation by NOX-generated ROS promotes survival following radiation. A) Western blot of shCON or shp22 cells (HK157) 5 days after radiation with 0 Gy or 10 Gy. p22phox, PTEN oxidation and total and phospho-Akt with actin used as a loading control. PTEN quantitation is of oxidized PTEN. Image is representative n>3 B) Changes in cell number were determined with a CCK-8 assay in shCON or shPTEN cells (HK157) 5 days after exposure to 0 Gy or 10 Gy with or without p22phox KD. n>3, *=*p*-value<0.01 as compared to shCON C) Changes in ROS levels were determined with CellROX Green in shCON or shPTEN cells (HK157) 5 days after exposure to 0 Gy or 10 Gy with or without p22phox KD. n>3, *=*p*-value<0.01 as compared to shCON

Finally, we wanted to determine if NOX inhibition sensitizes GBM cells to radiation *in vivo*. As expected, radiation decreased tumor burden in mice intracranially transplanted with PTEN functional GBM cells (shCON) (Fig. 7A-B). Interestingly, we found that knockdown of p22 alone was sufficient to decrease tumor burden. However, the combination of p22 knockdown and radiation resulted in mice with the least amount of tumor burden. When compared to shCON and irradiated mice, we found that knockdown of p22 significantly decreased tumor burden in mice suggesting that p22, and hence the NOX complex, promote resistance to radiation in GBM. Tumors were examined for p22 expression and Akt phosphorylation (given the methodology, it was not possible to examine PTEN oxidation in these cells) and found that Akt phosphorylation was increased following radiation (Fig. 7C) and this increase was lost in tumors lacking p22. Collectively, our data shows that NOX generates ROS following radiation, which oxidizes and activates PTEN resulting in increased Akt phosphorylation which in turn promotes proliferation. This suggests that inhibition of the NOX complex in PTEN functional tumors could improve radiation sensitivity in GBM.

**Figure 7:**
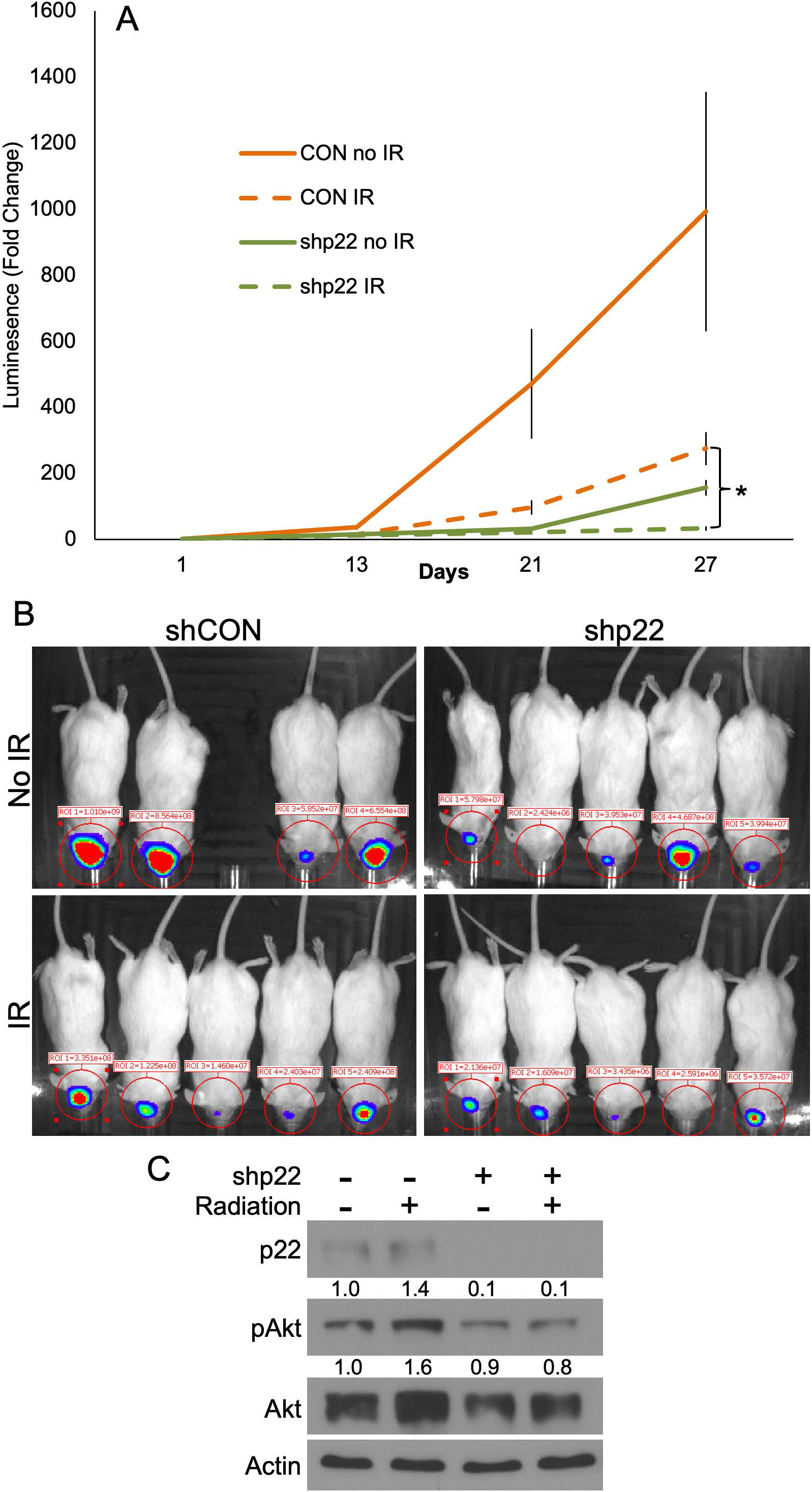
NOX promotes radio-resistance *in vivo*. A) Tumor volume was assessed using luminescence from shCON or shp22 GFP-FLUC xenograft intracranial tumors (HK408) at different time points following a single dose of radiation. *=*p*-value<0.01 as compared to shCON B) Images of different treatment groups on day 21 following radiation C) Western blot of shCON or shp22 non-perfused xenograft intracranial tumors (HK408) 28 days after radiation

## DISCUSSION

Here we report that low oxygen or radiation oxidizes PTEN resulting in its inactivation and subsequent activation of Akt. We also show that the NOX complex is activated following radiation and both hypoxia and radiation increase intracellular ROS levels through NOX. Finally, we found the inhibition of the NOX pathway sensitized GBM cells to radiation both *in vitro* and *in vivo*.

Reactive oxygen species are highly reactive chemicals formed from O_2_ and include, peroxides, super oxides, hydroxyl radicals and singlet oxygen. Exogenous sources of ROS include pollutants and radiation, while endogenous sources primarily come from cellular metabolism. Low levels of ROS are required for normal cellular function, while high levels of ROS promote apoptosis, autophagy, necrosis and ferroptosis. Recently, moderate increase in ROS levels have been shown to promote tumorigenesis, invasion and metastasis, without going high enough to induce the negative effects of ROS (Gong et al., 2022). Given the complexity of this homeostasis, we decided to examine how moderate increases in ROS from the microenvironment and treatment would affect GBM growth and response to therapy.

The PTEN protein is made of two primary domains, a tyrosine phosphatase domain (which contains the active site) and a C2 domain which binds it to the plasma membrane. In the presence of H_2_O_2_, a disulfide bond is formed between C71 and C124, shutting down the active site so the protein can no longer dephosphorylate PIP3 which results in prolonged activation of Akt (Zhang et al., 2020). In our hands, we found that PTEN was oxidized when cells were grown under biologically relevant conditions (like the low oxygen environment of tumors) and following a single treatment with radiation. This in turn correlated with an increase in Akt activation, cell numbers and radio-resistance. Treatment with an Akt inhibitor under hypoxic conditions still resulted in PTEN oxidation, however Akt activation and increased cell numbers were lost, suggesting that the PTEN/Akt pathway is required for the biological effects reported in this paper.

While some GBM express virtually no functional PTEN; due to deletion, mutation, or both, many express functional PTEN protein or only have PTEN mutations in one allele. However, the vast majority of tumors exhibit at least some activation of the PI3K pathway (Verhaak et al., 2010). While a number of different mechanisms can promote downstream Akt and mTOR activity, including crosstalk from the MAPK pathway, direct mutations in PI3 kinases and other relevant proteins (Ludwig and Kornblum, 2017), our data suggest that the hypoxic environment of the tumor may play a role in activation of this pathway, and thus antioxidant treatment may be an interesting direction of research—one that might seem counterintuitive at first glance. In fact, laboratory data has found that curcumin (a popular antioxidant) is able to sensitize glioblastoma cells to radiation through a variety of pathways (Zoi et al., 2022). However, our study would suggest that at least some residual PTEN activity would be required for a significant therapeutic effect. We do not yet know whether functional PTEN is required for the pro-tumorigenic role of ROS in other cancers.

One of the primary sources of endogenous ROS is the NADPH oxidase (NOX) complex of proteins found on cell membranes. Increased activation or expression of one or several NOX member proteins have been reported in multiple cancers corresponding with increased tumorgenicity and resistance to treatment (Mortezaee et al., 2019). Previous studies have demonstrated that activation of NOX results in enhanced neural stem cell self-renewal and enhanced neurogenesis in a PTEN dependent manner (Le Belle et al., 2011). Although it is unclear whether brain tumors are derived from neural stem cells, we reasoned that the shared properties of neural stem and glioblastoma stem cells might extend to a shared mechanism of ROS-induced proliferation. This, indeed, proved to be the case. In fact, our study goes beyond the notion that PTEN is required for the effects of NOX activation in glioma and neural stem cells, but that under physiological conditions in which NOX is activated, such as during hypoxia, PTEN is oxidized and inactivated, leading to the activation of Akt. Further studies will be required to determine whether there are distinct functional differences between the effects of NOX activation and outright PTEN deletion in GBM.

The induction of ROS is the main underlying mechanism of radiotherapy and several prior studies have found that inhibition of NOX increases sensitivity to radiation however the mechanism of action has yet to be elucidated (Teixeira et al., 2017). Here, we demonstrated that radiation induces NOX mRNA and activates NOX which, in turn, oxidizes PTEN, ultimately leading to downstream activation of Akt and enhanced cell production *in vitro* and enhanced tumor growth *in vivo*. These findings support the concept of NOX inhibition as an adjunct to radiation therapy in PTEN replete tumors. In fact, because of the importance of the NOX family of proteins, in several disorders, including cancer and stroke, a great deal of effort has gone into developing pharmacological NOX inhibitors. Recently the WHO approved a new stem, “naxib” which refers to **<u>NA</u>**DPH o**<u>x</u>**idase inh**<u>ib</u>**itor and thereby recognized NOX inhibitors as a new therapeutic class (Elbatreek et al., 2021). With the development of these and other drugs, it may eventually be possible to supplement radiation treatment for PTEN positive GBM patients with a NOX inhibitor and see improved outcomes.

There are several limitations to our study. First, we do not yet know whether the NOX system is truly expressed in GBM cells in human tumors *in situ*. While NOX2 and NOX4 are both expressed in GBM, review of single cell studies reveals limited expression of the mRNA for NOX components within tumor cells (Darmanis et al., 2017). However, we do not know whether this limited expression is sufficient to play a direct role in tumor biology in humans or whether NOX is induced by radiation in human GBM cells undergoing therapeutic radiation. It is possible that NOX is upregulated by the act of growing them in glioma stem cell-enriched gliomasphere cultures and that this upregulation is maintained during *in vivo* growth in xenografts. It is possible that human brain tumors, including those undergoing radiation, receive ROS via non-cell autonomous activation of other cells in the microenvironment, including myeloid and vascular cells (Bekhet and Eid, 2021), both of which express abundant mRNA for NOX components (Darmanis et al., 2017).

Another limitation of our current study is in the dose and schedule of radiation used. We chose a single, high dose of radiation, rather than a fractionated schedule as is administered to people (Ziu et al., 2020). Murine xenografts and human tumors *in situ* are of vastly different sizes (Ma et al., 2020; Rutter et al., 2017). A radiation dose that kills a large percentage of cells may leave tens of millions of cells leftover in a human tumor, and only a few cells left in a murine tumor, thus making it unlikely to be able to observe tumor growth and regrowth following radiation over the lifespan of a mouse. Future preclinical studies using either large animal models, or alternate dosing schedules may be required prior to therapeutic human trials with NOX inhibitors and GBM radiation. Despite these limitations, our study provides a strong basis for pursuing NOX inhibition in PTEN-expressing GBM cells as a possible adjunct to radiation therapy.

## INNOVATION

While other studies have implicated NOX function in GBM models, these studies demonstrate NOX activation and function under physiological hypoxia and following radiation in GBM, two conditions that are seen in patients. NOX plays an important role in a PTEN-expressing GBM model system, but not in PTEN-non-functional systems and provide a potential, patient-specific therapeutic opportunity.

## MATERIALS AND METHODS

### Tissue culture

The isolation and propagation of the primary glioma stem cell containing cultures (gliomaspheres) used for this study was described previously (Laks et al., 2016). The prior manuscript also describes the patient characteristics for each line used. Briefly, to culture the cells, on the day of resection samples were digested with TryplE and further dissociated mechanically. Acellular debris was removed, and remaining cells were incubated in gliomasphere media (DMEM/F12 supplemented with B27, Penicillin/ Streptomycin, heparin, EGF and bFGF) for several days until spheres began to form. Frozen stocks were made at passage 5 to maintain cells at low passage. Cell cultures were maintained as previously described (Laks et al., 2009).

### Drug treatment

Cells were plated according to their assay at a density of 100,000 cells/mL and allowed to settle for 24 hours. Following that time, cells were treated with a single dose of the following drugs and most experiments were run after 5 days. Hydrogen Peroxide (H_2_O_2_), N-acetylcysteine (NAC), LY-294002 (LY), or Apocynin (APO).

### Immunofluorescence staining and quantitation

For immunostaining, cells were first fixed in 4% PFA for 15 minutes. Following fixation, cells were washed with PBS, permeabilized with PBS with 0.1% Triton X-100 for nuclear staining and blocked in PBS with 2% BSA at room temperature for 30 minutes. Cells were then incubated with the indicated primary antibodies: phospho-yH2AX (1:100, Cell Signaling, # 9718), NOX2 (1:100, Thermo Fisher Scientific, PA5-79118), NOX4 (1:100, R&D systems, MAB8158), and CYBA/P22 (1:100, Thermo Fisher Scientific, PA5-71642) overnight at 4°C. Cells were washed with PBS, and incubated with species-appropriate goat/donkey secondary antibodies coupled to AlexaFluor dye 568 (Invitrogen) and Hoechst dye for nuclear staining for 2 hrs at RT. Stained cells were imaged using EVOS Cell Imaging systems microscope (Thermo Fisher Scientific), and quantification was performed using Image J. For quantitation, phospho-yH2AX+ and Hoechst+ cells were counted in at least 5 images per condition, and data is represented as mean ± SD in the graphs.

### Western blots

Proteins from the cell lines were extracted in RIPA buffer and quantitated by Bio-Rad protein assay. Equal amounts of total proteins were loaded on SDS-PAGE gels, transferred onto a nitrocellulose membrane, and probed with primary antibodies (anti-PTEN [Cell Signaling], anti-actin [Abcam], anti-p22phox [Abcam], anti-Akt [Cell Signaling] and anti-phospho-Akt T308 [Cell Signaling]) overnight at 4°C. HRP conjugated secondary antibodies were incubated for 1 hour at room temperature. Proteins were visualized by chemiluminescence as recommended by the manufacturer (Thermo Fisher Scientific). PTEN oxidation was assessed according to (Delgado-Esteban et al., 2007).

### Animal strains, intracranial Xenotransplantation and Optical imaging

All animal studies were performed according to approved protocols by the institutional animal care and use committee at UCLA. Studies did not discriminate sex, and both male and females were used. 8-to 12-week-old NOD-SCID gamma null (NSG) mice were used to generate tumors from a patient-derived GBM line HK408. 5×10^4^ tumor cells containing a firefly-luciferase (FLUC)-GFP lentiviral construct were injected intracranially into the neostriatum in mice. Tumor growth was monitored at 2, 3 and 4 weeks after transplantation using IVIS Lumina II bioluminescence imaging at the Crump Institute for Molecular Imaging at UCLA. Mice were anesthetized by inhalation of isoflurane. Intraperitoneal injection of 100uL of luciferin (30mg/ml) was followed by 10 minutes of live uptake to interact with the luciferase expressing FLUC-GFP tumor cells and produce bioluminescence. The IVIS Lumina 2 imaging system (Caliper life sciences) was utilized to photograph the mice, and images were overlaid with a color scale of a region of interest representing total flux (photon/second) and quantified with the Living Image software package (Xenogen).

### ROS Measurement

Cells were grown at a density of 100,000 cell/mL and allowed to grow for the appropriate time following treatment. On the day of experiment, spheres were dissociated with Accumax (Innovative Cell Technologies) and live cells were counted using a Countess Automated Cell Counter (Life Technologies), with 0.4% trypan blue exclusion. 50,000 cells were treated with CellROX Green (Thermo Fisher) or MitoSOX (Thermo Fisher) according to the manufacturers protocol. Cells were then counterstained with Hoechst 33342 (Thermo-Fisher) to use as a loading control for calculations. Cells were loaded into a 96-well black well plate and read on a Varioskan Lux multimode plate reader and fluorescence was recorded at the appropriate wavelengths.

### Growth Assay

Cells were grown at a density of 100,000 cell/mL and allowed to grow for the appropriate time following treatment. On the day of experiment, spheres were treated with Cell Counting Kit -8 (Dojindo) according to the manufacturers protocol. Results were read with a microplate reader on a Varioskan Lux multimode plate reader and absorbance was recorded at 450nm.

### Irradiation of glioma cultures and orthotopic tumor xenografts

Cultured cells were irradiated at room temperature using an experimental X-ray irradiator (Gulmay Medical Inc. Atlanta, GA) at a dose rate of 5.519 Gy/min for the time required to apply a prescribed dose. The X-ray beam was operated at 300 kV and hardened using a 4mm Be, a 3mm Al, and a 1.5mm Cu filter and calibrated using NIST-traceable dosimetry. Tumor-bearing mice were irradiated at a single dose of 10 Gy using an image-guided small animal irradiator (X-RAD SmART, Precision X-Ray, North Branford, CT) with an integrated cone beam CT (60 kVp, 1 mA) and a bioluminescence-imaging unit as described previously (Bhat et al., 2020). Individual treatment plans were calculated for each animal using the SmART-Plan treatment planning software (Precision X-Ray). Radiation treatment was applied using a 5×5mm collimator from a lateral field.

### qRT-PCR

RNA was isolated using TRIzol (Gibco) and 1.5 µg RNA was converted to cDNA by reverse transcription. qRT-PCR was performed after addition of Power SYBR Master Mix (Applied Biosystems) on an ABI PRISM 7700 sequence detection system (Applied Biosystems).

### shRNA

Lentiviral mediated shRNA knockdowns of *PTEN* and *p22-phox* were performed using constructs from the Dharmacon-Harmon library. Four shPTEN constructs and five shp22-phox constructs were tested, and one of each was chosen following examination of knockdown efficiency and resulting phenotype. Briefly, virus was made using HEK-293T cells transfected with package, envelope and the shRNA construct using lipofectamine (Thermo-Fisher) in the absence of serum or antibiotics. Conditioned media from the cells was collected three days later and added to the GBM cell line of interest.

### Statistics

For comparison of small groups, we used a cutoff of *p* < .05 to distinguish significant differences. Statistics for comparing cell numbers, ROS levels, mRNA and γH2AX staining were done with ANOVA analysis and posthoc t-tests.

## AUTHOR CONTRIBUTIONS

K.L., S.D.M, M.C. and J.S. designed, performed, and analyzed the experiments. J.L.B., V.E., F.J., and H.K. conceived and conceptualized the study. K.L., S.D.M, M.C., and H.K. wrote the article. All authors reviewed the article.

## FIGURE LEGENDS

**Supplementary Figure 1:**
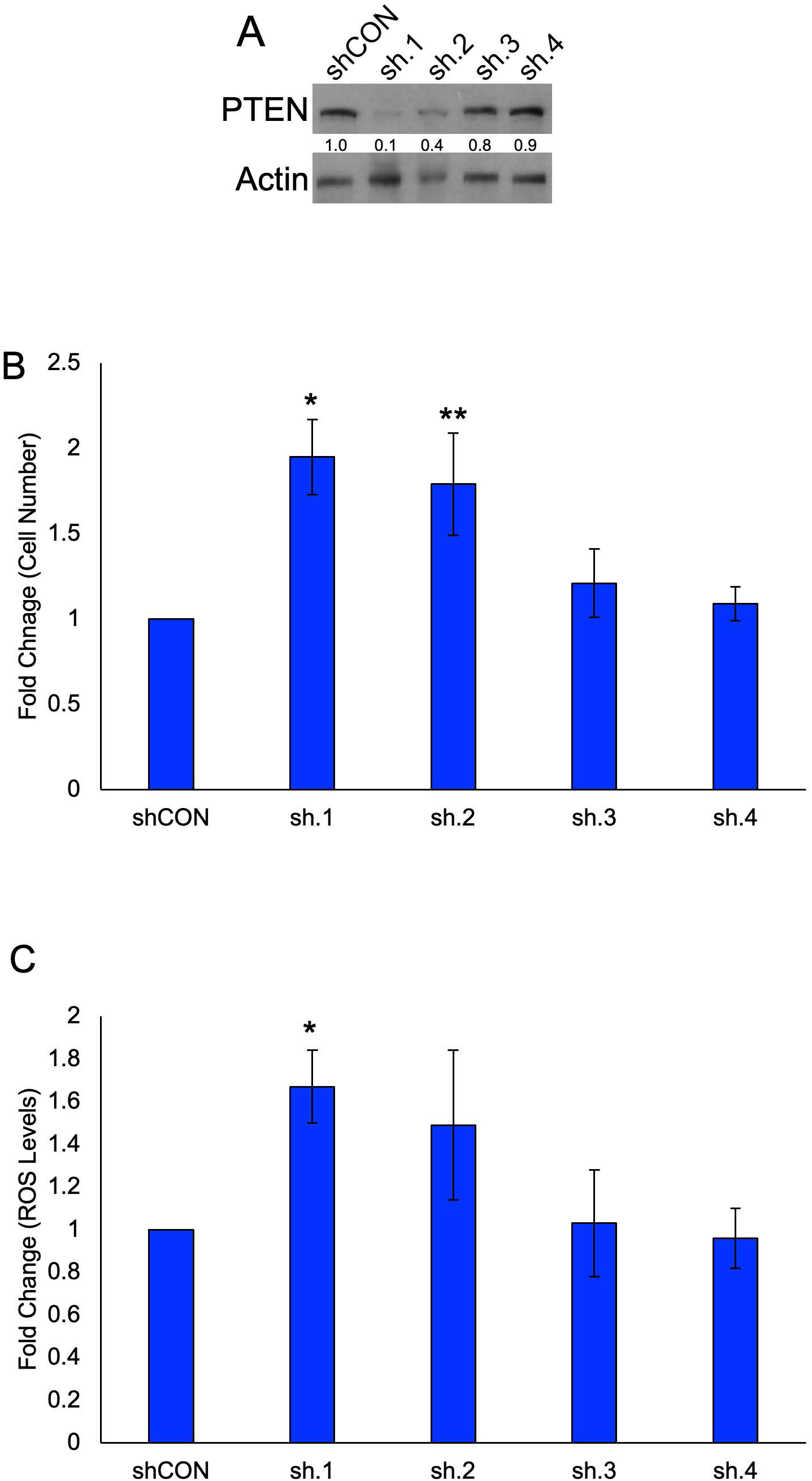
Physiological effects of PTEN knockdown. A) Western blot of HK157 cells following knockdown of PTEN with 4 different clones. PTEN with actin used as a loading control. Image is representative n>3 B) Changes in cell number were determined with a CCK-8 assay following knockdown for 5 days in HK157 with either shCON or shPTEN. n>3, *=*p*-value<0.01 as compared to shCON, **=*p*-value<0.05 as compared to shCON. C) Changes in ROS levels were determined with CellROX Green following knockdown for 5 days in HK157 with either shCON or shPTEN. n>3, **=*p*-value<0.05 as compared to shCON.

**Supplementary Figure 2:**
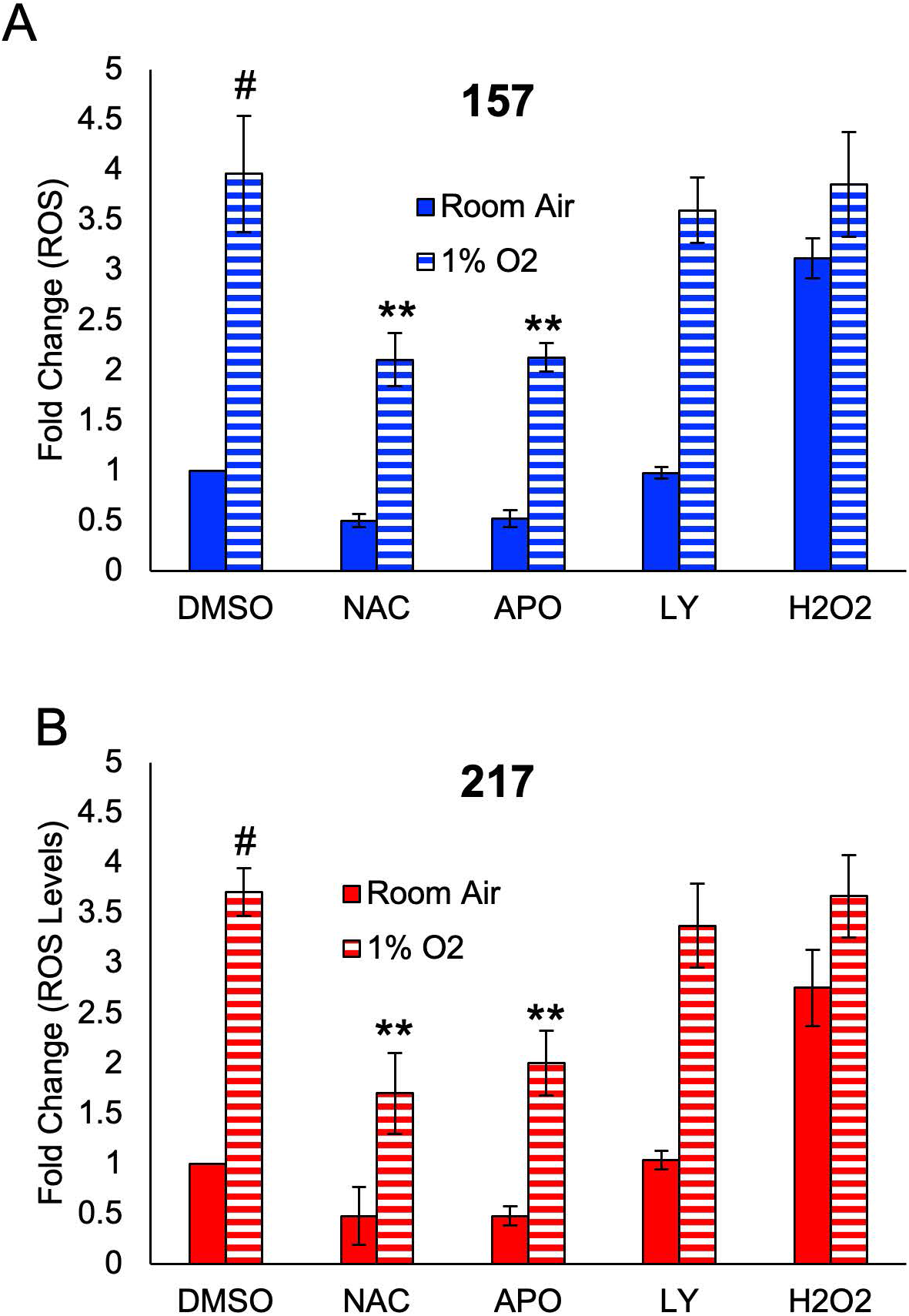
Hypoxia increases ROS levels. A) Changes in ROS levels were determined with CellROX Green in a PTEN functional line (HK157) grown under room air or 1% O_2_ for 5 days and treated with NAC (1mM), APO (100uM), LY (20uM), or H_2_O2 (10uM). n>3, #=*p*-value<0.01 as compared to room air, **=*p*-value<0.05 as compared to DMSO, 1% 02. B) Changes in ROS levels were determined with CellROX Green in a PTEN non-functional line (HK217) grown under room air or 1% O_2_ for 5 days and treated with NAC (1mM), APO (100uM), LY (20uM), or H_2_O2 (10uM). n>3, #=*p*-value<0.01 as compared to room air, **=*p*-value<0.05 as compared to DMSO, 1% 02.

**Supplementary Figure 3:**
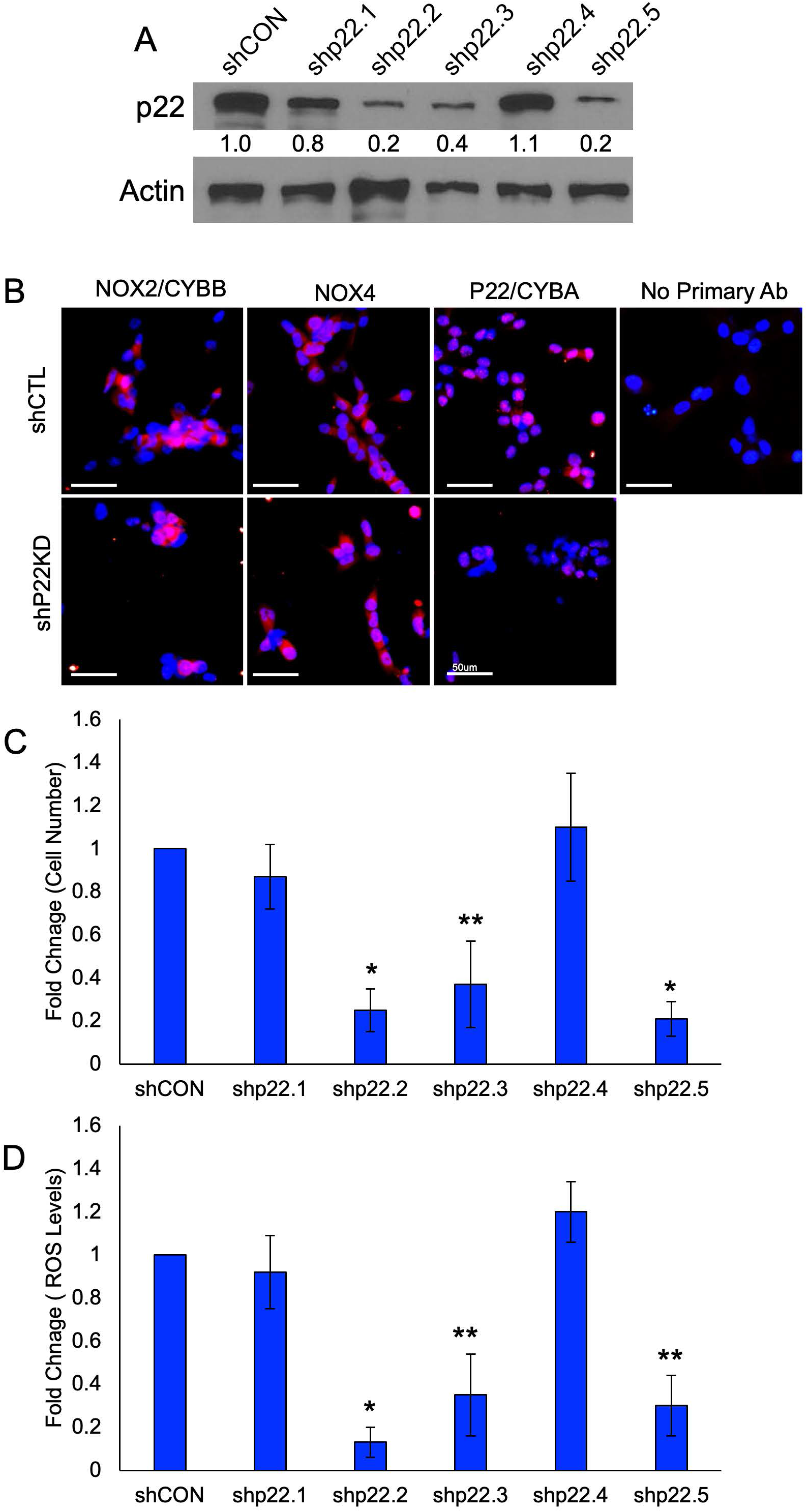
Knockdown of p22phox. A) Western blot of HK157 cells following knockdown of p22phox with 5 different clones. p22 with actin used as a loading control. Image is representative n>3 B) Representative immunocytochemistry images of CYBA, CYBAA and NOX4 following shCON or shp22 treatment in HK408 cells. C) Changes in cell number were determined with a CCK-8 assay following knockdown for 5 days in HK157 with either shCON or sh22. n>3, *=*p*-value<0.01 as compared to shCON, **=*p*-value<0.05 as compared to shCON. D) Changes in ROS levels were determined with CellROX Green following knockdown for 5 days in HK157 with either shCON or sh22. n>3, **=*p*-value<0.05 as compared to shCON.

## REFERENCES

Bekhet OH and Eid ME. The Interplay between Reactive Oxygen Species and Antioxidants in Cancer Progression and Therapy: A Narrative Review. Transl Cancer Res 2021;10(9):4196–4206; doi: 10.21037/tcr-21-629.

Bhat K, Saki M, Vlashi E, et al. The Dopamine Receptor Antagonist Trifluoperazine Prevents Phenotype Conversion and Improves Survival in Mouse Models of Glioblastoma. Proc Natl Acad Sci U S A 2020;117(20):11085–11096; doi: 10.1073/pnas.1920154117.

Bowman RL, Wang Q, Carro A, et al. GlioVis Data Portal for Visualization and Analysis of Brain Tumor Expression Datasets. Neuro-Oncol 2017;19(1):139–141; doi: 10.1093/neuonc/now247.

Cao W, Zhou Q, Wang H, et al. Hypoxia Promotes Glioma Stem Cell Proliferation by Enhancing the 14-3-3β Expression via the PI3K Pathway. J Immunol Res 2022;2022:5799776; doi: 10.1155/2022/5799776.

Chen Y, Li Y, Huang L, et al. Antioxidative Stress: Inhibiting Reactive Oxygen Species Production as a Cause of Radioresistance and Chemoresistance. Oxid Med Cell Longev 2021;2021:6620306; doi: 10.1155/2021/6620306.

Darmanis S, Sloan SA, Croote D, et al. Single-Cell RNA-Seq Analysis of Infiltrating Neoplastic Cells at the Migrating Front of Human Glioblastoma. Cell Rep 2017;21(5):1399–1410; doi: 10.1016/j.celrep.2017.10.030.

Delgado-Esteban M, Martin-Zanca D, Andres-Martin L, et al. Inhibition of PTEN by Peroxynitrite Activates the Phosphoinositide-3-Kinase/Akt Neuroprotective Signaling Pathway. J Neurochem 2007;102(1):194–205; doi: 10.1111/j.1471-4159.2007.04450.x.

Elbatreek MH, Mucke H and Schmidt HHHW. NOX Inhibitors: From Bench to Naxibs to Bedside. Handb Exp Pharmacol 2021;264:145–168; doi: 10.1007/164_2020_387.

Gong S, Wang S and Shao M. NADPH Oxidase 4: A Potential Therapeutic Target of Malignancy. Front Cell Dev Biol 2022;10:884412; doi: 10.3389/fcell.2022.884412.

Groszer M, Erickson R, Scripture-Adams DD, et al. Negative Regulation of Neural Stem/Progenitor Cell Proliferation by the Pten Tumor Suppressor Gene in Vivo. Science 2001;294(5549):2186–2189; doi: 10.1126/science.1065518.

Groszer M, Erickson R, Scripture-Adams DD, et al. PTEN Negatively Regulates Neural Stem Cell Self-Renewal by Modulating G0-G1 Cell Cycle Entry. Proc Natl Acad Sci U S A 2006;103(1):111–116; doi: 10.1073/pnas.0509939103.

Hemmati HD, Nakano I, Lazareff JA, et al. Cancerous Stem Cells Can Arise from Pediatric Brain Tumors. Proc Natl Acad Sci U S A 2003;100(25):15178–15183; doi: 10.1073/pnas.2036535100.

Laks DR, Crisman TJ, Shih MYS, et al. Large-Scale Assessment of the Gliomasphere Model System. Neuro-Oncol 2016;18(10):1367–1378; doi: 10.1093/neuonc/now045.

Laks DR, Masterman-Smith M, Visnyei K, et al. Neurosphere Formation Is an Independent Predictor of Clinical Outcome in Malignant Glioma. Stem Cells Dayt Ohio 2009;27(4):980–987; doi: 10.1002/stem.15.

Le Belle JE, Orozco NM, Paucar AA, et al. Proliferative Neural Stem Cells Have High Endogenous ROS Levels That Regulate Self-Renewal and Neurogenesis in a PI3K/Akt-Dependant Manner. Cell Stem Cell 2011;8(1):59–71; doi: 10.1016/j.stem.2010.11.028.

Le Belle JE, Sperry J, Ngo A, et al. Maternal Inflammation Contributes to Brain Overgrowth and Autism-Associated Behaviors through Altered Redox Signaling in Stem and Progenitor Cells. Stem Cell Rep 2014;3(5):725–734; doi: 10.1016/j.stemcr.2014.09.004.

Ludwig K and Kornblum HI. Molecular Markers in Glioma. J Neurooncol 2017;134(3):505–512; doi: 10.1007/s11060-017-2379-y.

Ma Z, Niu B, Phan TA, et al. Stochastic Growth Pattern of Untreated Human Glioblastomas Predicts the Survival Time for Patients. Sci Rep 2020;10(1):6642; doi: 10.1038/s41598-020-63394-w.

McKetney J, Runde RM, Hebert AS, et al. Proteomic Atlas of the Human Brain in Alzheimer’s Disease. J Proteome Res 2019;18(3):1380–1391; doi: 10.1021/acs.jproteome.9b00004.

Mortezaee K, Goradel NH, Amini P, et al. NADPH Oxidase as a Target for Modulation of Radiation Response; Implications to Carcinogenesis and Radiotherapy. Curr Mol Pharmacol 2019;12(1):50–60; doi: 10.2174/1874467211666181010154709.

Rodriguez SMB, Staicu G-A, Sevastre A-S, et al. Glioblastoma Stem Cells-Useful Tools in the Battle against Cancer. Int J Mol Sci 2022;23(9):4602; doi: 10.3390/ijms23094602.

Rutter EM, Stepien TL, Anderies BJ, et al. Mathematical Analysis of Glioma Growth in a Murine Model. Sci Rep 2017;7(1):2508; doi: 10.1038/s41598-017-02462-0.

Su X, Yang Y, Guo C, et al. NOX4-Derived ROS Mediates TGF-Β1-Induced Metabolic Reprogramming during Epithelial-Mesenchymal Transition through the PI3K/AKT/HIF-1α Pathway in Glioblastoma. Oxid Med Cell Longev 2021;2021:5549047; doi: 10.1155/2021/5549047.

Teixeira G, Szyndralewiez C, Molango S, et al. Therapeutic Potential of NADPH Oxidase 1/4 Inhibitors. Br J Pharmacol 2017;174(12):1647–1669; doi: 10.1111/bph.13532.

Verhaak RGW, Hoadley KA, Purdom E, et al. Integrated Genomic Analysis Identifies Clinically Relevant Subtypes of Glioblastoma Characterized by Abnormalities in PDGFRA, IDH1, EGFR, and NF1. Cancer Cell 2010;17(1):98–110; doi: 10.1016/j.ccr.2009.12.020.

Vermot A, Petit-Härtlein I, Smith SME, et al. NADPH Oxidases (NOX): An Overview from Discovery, Molecular Mechanisms to Physiology and Pathology. Antioxid Basel Switz 2021;10(6):890; doi: 10.3390/antiox10060890.

Vilar JB, Christmann M and Tomicic MT. Alterations in Molecular Profiles Affecting Glioblastoma Resistance to Radiochemotherapy: Where Does the Good Go? Cancers 2022;14(10):2416; doi: 10.3390/cancers14102416.

Zhang Y, Park J, Han S-J, et al. Redox Regulation of Tumor Suppressor PTEN in Cell Signaling. Redox Biol 2020;34:101553; doi: 10.1016/j.redox.2020.101553.

Ziu M, Kim BYS, Jiang W, et al. The Role of Radiation Therapy in Treatment of Adults with Newly Diagnosed Glioblastoma Multiforme: A Systematic Review and Evidence-Based Clinical Practice Guideline Update. J Neurooncol 2020;150(2):215–267; doi: 10.1007/s11060-020-03612-7.

Zoi V, Galani V, Tsekeris P, et al. Radiosensitization and Radioprotection by Curcumin in Glioblastoma and Other Cancers. Biomedicines 2022;10(2):312; doi: 10.3390/biomedicines10020312.

